# Composition, structure and robustness of Lichen guilds

**DOI:** 10.1101/2022.10.30.514411

**Authors:** Salva Duran-Nebreda, Sergi Valverde

## Abstract

Symbiosis is a major engine of evolutionary innovation underlying the many extant complex organisms. Lichens are a paradigmatic example that offers a unique perspective on the role of symbiosis in ecological success and evolutionary diversification. Lichen studies have produced a wealth of information regarding the importance of symbiosis in many different species, but they frequently focus on a few species, limiting our understanding of large-scale phenomena such as guilds. Guilds are groupings of lichens that assist each other’s proliferation and are intimately linked by a shared set of photobionts, constituting an extensive network of relationships. To characterize the network of lichen symbionts, we used a large data set (*n* = 206 publications) of natural photobiont-mycobiont associations. The entire lichen network was found to be modular, but this organization does not replicate taxonomic information in the data set, prompting a reconsideration of lichen guild structure and composition. The characteristic scale of effective information reveals that the major lichen guilds are better represented as clusters with several substructures rather than as monolithic communities. Heterogeneous guild structure fosters robustness, with keystone species functioning as bridges between guilds and whose extinction would endanger global stability.

## Introduction

Lichens are symbiotic organisms composed of a fungus (mycobiont), one or more photosynthetic partners (photobionts, typically algae or cyanobacteria see figure 1a) and other microbial species^1,2^. In the wider ecological context, lichens provide several services that are essential for ecosystem functioning: from weathering of rocks increasing the bioavailability of minerals^3^ to carbon and nitrogen fixation^4^. By virtue of the wildly different metabolisms of photobionts and mycobionts, lichenization provides new biological traits that can enable both partners to colonize a wider range of environments^5,6^, including extreme^3,7^, polluted^8,9^ or anthropogenic ecosystems^10^. In consequence, lichens are more than the sum of their constituent symbionts, emphasizing the relevance of non-fraternal organismality as a source of evolutionary innovation.^11^.

**Figure 1.**
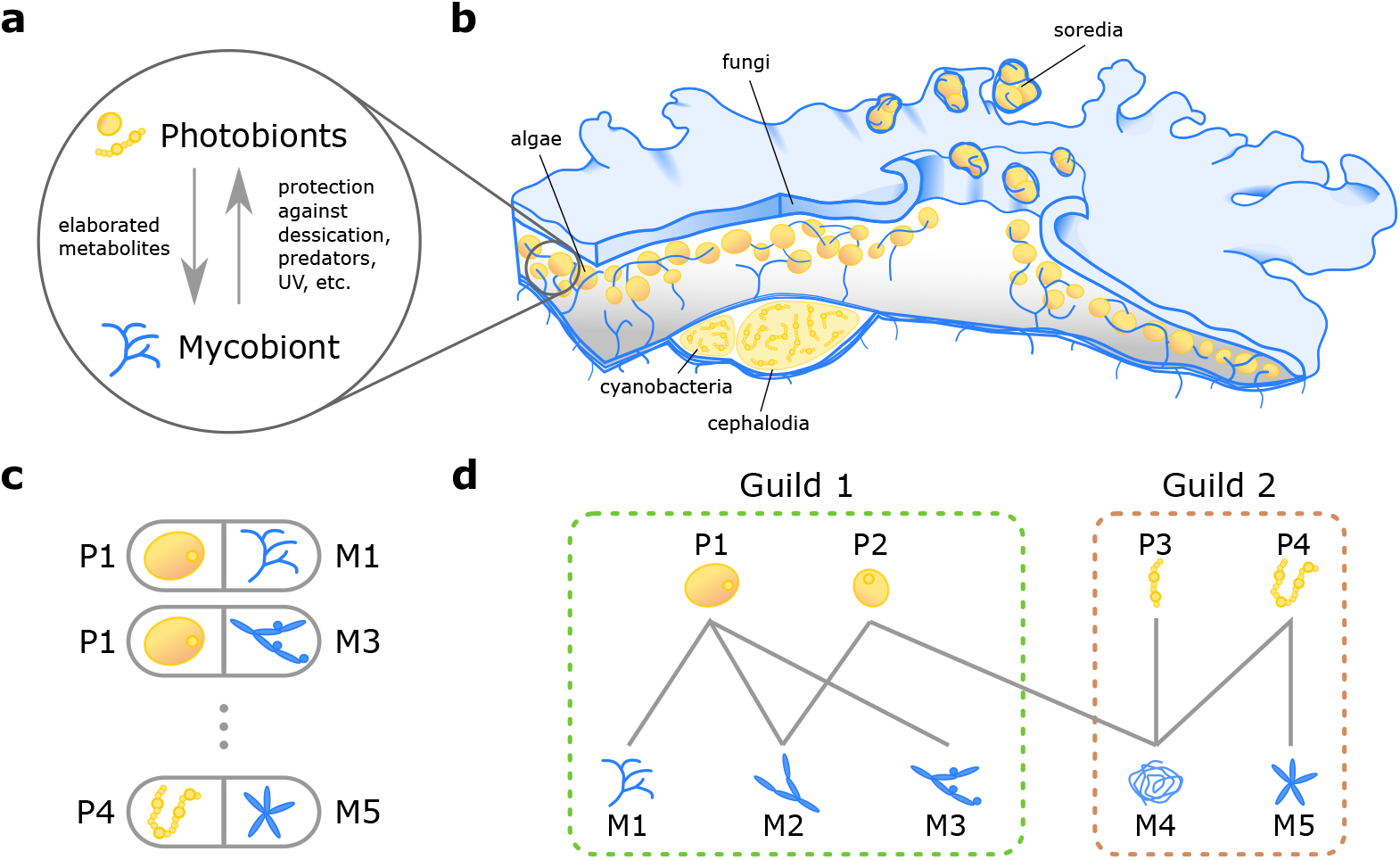
A network perspective on lichen symbioses. (a) schematic representation of lichen symbiosis at the ecological level. Photobionts (yellow) provide energy fixation though elaborated metabolites while the mycobiont (blue) comprises the majority of the lichen body and provides protection to various environmental challenges. (b) Example lichen body, including distinct organs to segregate algal and cyanobacterial photobionts (e.g. cephalodia). Here, lichen reproduction takes place asexually though soredia, which include both the algal (yellow) and fungal (blue) partners in each propagule. (c) Each individual lichen species (grey boxes) is composed by a mycobiont (*M*_*j*_) and one or more photobionts (*P*_*i*_). (d) The set of associations among photobionts and mycobionts defines a bipartite network, its structural analysis can reveal the presence of mesoscale structures, like guilds, related to the underlying ecological relations and evolutionary history.

Symbiosis is a natural hot spot for functional diversity^12,13^, the evolutionary potential of a lichen symbiont must take into account the extraordinary repertoire of traits gained by acquiring a new partner. However, it is widely acknowledged that specificity of the mycobiont-photobiont association as well as the spatial distribution of their component species largely drives the formation of new lichen organisms^6,14–16^. Indeed, many symbionts are quite stringent in the partnerships they form, which are often determined by biophysical gradients^17–22^. It has also been suggested that highly specific mycobionts are typically restricted to few environments and display limited ecological range^23,24^. Conversely, mycobionts capable of colonizing different habitats often are less stringent in their partnerships, interacting with various available photobionts^12,16^.

Collectively, the set of natural associations between photobionts and mycobionts defines a network of symbionts^25,26^, where each interaction corresponds to a singular lichen species. It has been proposed that within this network lichen species organize in communities known as photobiont-mediated guilds^27^. Guilds are groups of species that are ecologically connected by sharing one or more photobionts^27,28^. Fungal species in a guild can benefit each other by propagating a common set of partners, driving the establishment of connected species into new or marginal habitats^29^, while competing for space and resources^30^.

Much of our current understanding on photobiont-mycobiont partnerships comes from studies involving a few species or geographical areas^16,17,21,22,31–36^. However, mycobiont-photobiont partnerships do not happen in isolation, they are part of a larger web of interactions supporting the assembly and maintenance of communities, and understanding them requires a systemic approach. Recent studies involving small sets of species and their interactions have revealed diverging structural patterns: from distinct clusters of strongly interacting species in *Peltigera* lichens^36^ to an embedded specialist-generalist structure in *Nephroma*^31^. Different explanatory causes have been proposed for the observed nested and modular patterns, including evolutionary constraints^36^ as well as the variations in the nature of the underlying ecological interactions^31^ (i.e. mutualistic vs. non-mutualistic relations). The general organization of photobiont-mycobiont associations and its connection to guilds remains largely unknown.

Here, we address these open questions by reconstructing the photobiont-mycobiont network, aggregating decades of research using Sanders and Masumoto’s meta-study^37^. This network records many observations (*n* = 206 publications) about individual species, but it also highlights other critical scales that operate beyond the species level, such as photobiont-mediated guilds. We combine several network metrics in order to understand the topological signature of guilds. Our findings show that taxonomy alone cannot comprehensively recapitulate guild topology. Network modularity, in particular, does not completely predict species composition of guilds. Effective information is a newly suggested metric that indicates a unique, multiscale pattern of structural diversity within guilds. The heterogeneous structure uncovered by effective information fosters robustness, with keystone species functioning as bridges across guilds and whose removal promotes network fragmentation and drives global instability.

## Results

### Defining the global photobiont-mycobiont association network (PMAN)

Previous research has focused on lichenization among few species of mycobionts and photobionts, providing an insufficient understanding of symbiotic interactions at largest scales. For example, symbiotic ties could be have been shaped by the presence of additional interactions in a larger community. In this context, network techniques have been particularly successful in the study of mutualistic^38^ and antagonistic^39^ networks from small to large spatio-temporal scales^40^. Here, we recreate the largest network of lichen symbionts to date using the full data set of symbiont pairings assembled in a recent meta-study by Sanders and Masumoto^37^. This system belongs to the general class of bipartite networks, which have links between nodes of different types. Figure 2 displays the photobiont-mycobiont association network (PMAN) representing this dataset, where blue and yellow nodes correspond to mycobionts and photobionts respectively.

**Figure 2.**
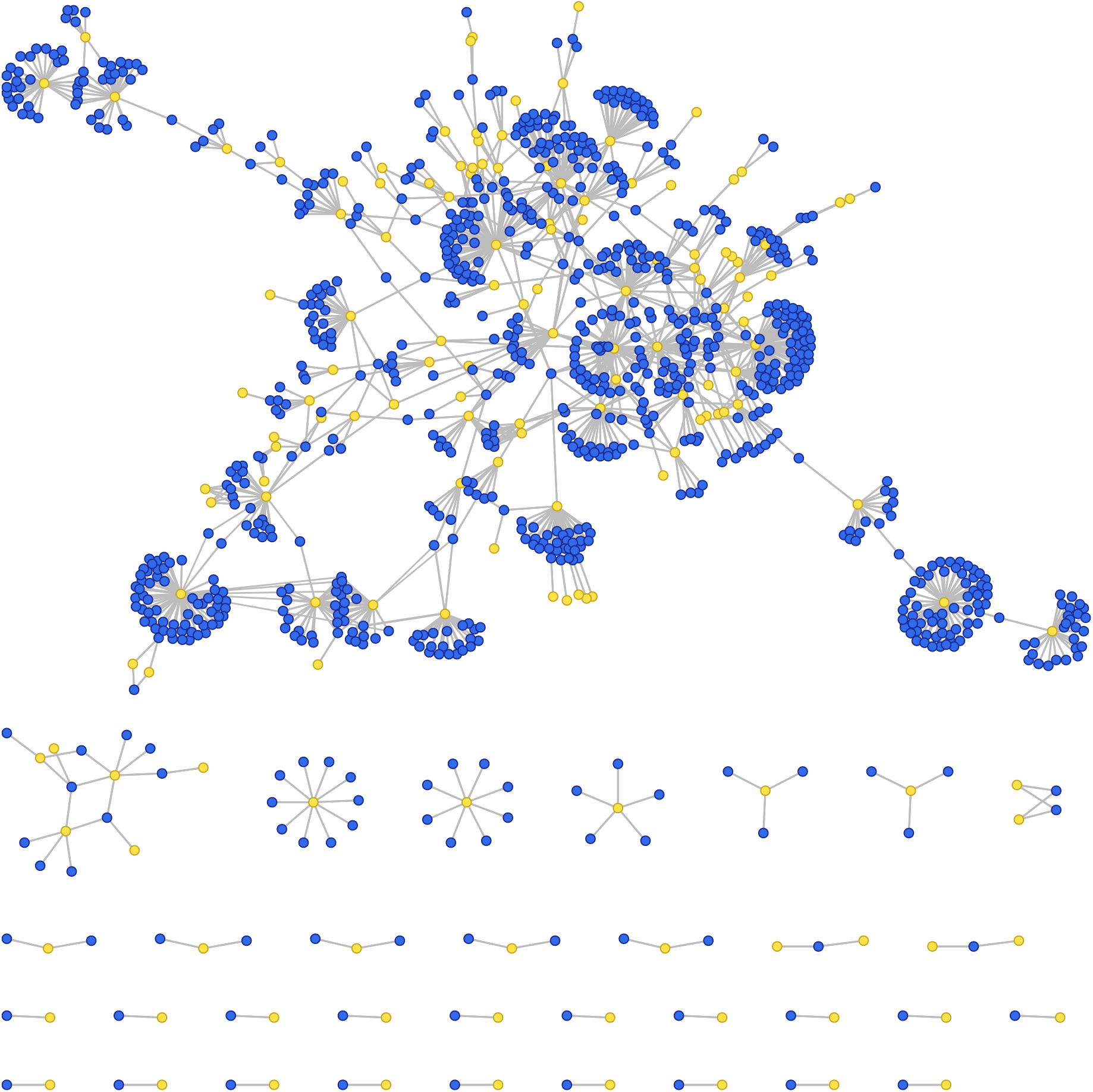
Photobiont-Mycobiont Association Network. Bipartite network representation of the full dataset analysed in our study. Interactions in this network involve two different types of nodes: blue nodes correspond to mycobionts, and yellow nodes indicate photobiont species. The network consists of 34 isolated components, the largest of which has average degree ⟨*k*⟩ = 2.524 (see Table 1). Network layout was automatically generated with the FMMM algorithm.

### The photobiont-mycobiont network is modular and not nested

Many species interaction networks are classified as nested or modular. A modular network is made up of sparsely linked clusters of dense subgraphs^38,41,42^. These communities or modules may arise due to evolutionary and environmental constraints^43^, or they may represent common functional qualities, such as groupings of closely related proteins implicated in cell communication. The other common structural feature of ecological networks is nestedness^44,45^. Nestedness relates to the hierarchical organisation in the network where nodes display a tendency to interact preferentially with subsets of partners of better-connected nodes^46^. This concept is particularly important in ecological network studies focusing on the spatial distribution of species, their interactions, and degree of individual specialisation. Modularity and nestedness are conceptually distinct structures that are negatively correlated; for example, a network with a high degree of modularity is typically associated with a low degree of nestedness. When compared to null models, however, certain empirical networks may exhibit both patterns^44^, suggesting the possibility of coexisting nestedness and modularity^39^.

**Table 1.** General properties of the photobiont-mycobiont network.

| General Properties | Definition | Value |
| --- | --- | --- |
| P | Number of Photobionts | 156 |
| M | Number of Mycobionts | 926 |
| S=P+M | Number of Species | 1082 |
| I | Number of Interactions | 1311 |
| $C=I/(P \times M)$ | Connectance | 0.00906 |
| $\langle k \rangle$ | Mean degree | 2.4196 |
| $\langle k_P \rangle$ | Mean Photobiont degree | 8.3910 |
| $Max(k_{Pi})$ | Max Photobiont degree | 75 |
| $\langle k_M \rangle$ | Mean Mycobiont degree | 1.4136 |
| $Max(k_{Mi})$ | Max Mycobiont degree | 9 |

To validate the presence of structural patterns, we compared the photobiont-mycobiont network to a bootstrap model that maintains the entire degree sequence for each compartment^42^ (see Methods). This gives a negative control that can help us determine the significance of the structures under consideration. Figure 3a shows the distribution of modularity values for an ensemble of bootstrap randomizations (histogram with shaded region) compared to the real data set average modularity (dashed vertical red line). This suggests that the lichen network is highly modular, more than the expected value for the null models (*p <* 10^−5^), consistently with a significant decrease in nestedness (*p <* 10^−5^, see Table 2). Following standard analyses of data completeness^47,48^, we studied the modularity and nestedness of subsamples using half the dataset, i.e. real networks and their randomizations with half the number of observed network links (see Fig 8). We discovered that modularity and nestedness are maintained in both circumstances, implying that the presented patterns are robust and independent of sampling depth.

**Table 2.** Structural properties of the photobiont-mycobiont network. From left to right, mean betweenness centrality for photobionts and mycobionts (*BC*_*P*_ and *BC*_*M*_ respectively), modularity (*Q*_*B*_) and whole nestedness (*NODF*). Null model metrics were calculated from 100 independent realizations of the network. P-values were obtained from t-test statistics in normally distributed variables (*Q*_*B*_, *NODF*) or Mann-Whitney U rank non-parametric test in non-normally distributed data (*BC*_*P*_ and *BC*_*M*_).

|  | Centrality |  |  |  | Modularity |  | Nestedness |  |
| --- | --- | --- | --- | --- | --- | --- | --- | --- |
| | $\langle BC_P \rangle$ | p-value | $\langle BC_M \rangle$ | p-value | $\langle Q_B \rangle$ | p-value | $\langle NODF \rangle$ | p-value |
| Data | 0.0188 | - | 0.0026 | - | 0.722 | - | 1.879 | - |
| Bootstrap | 0.01254 | 0.3435 | 0.0014 | 0.1848 | 0.691 | $< 10^{-5}$ | 2.419 | $< 10^{-5}$ |

**Figure 3.**
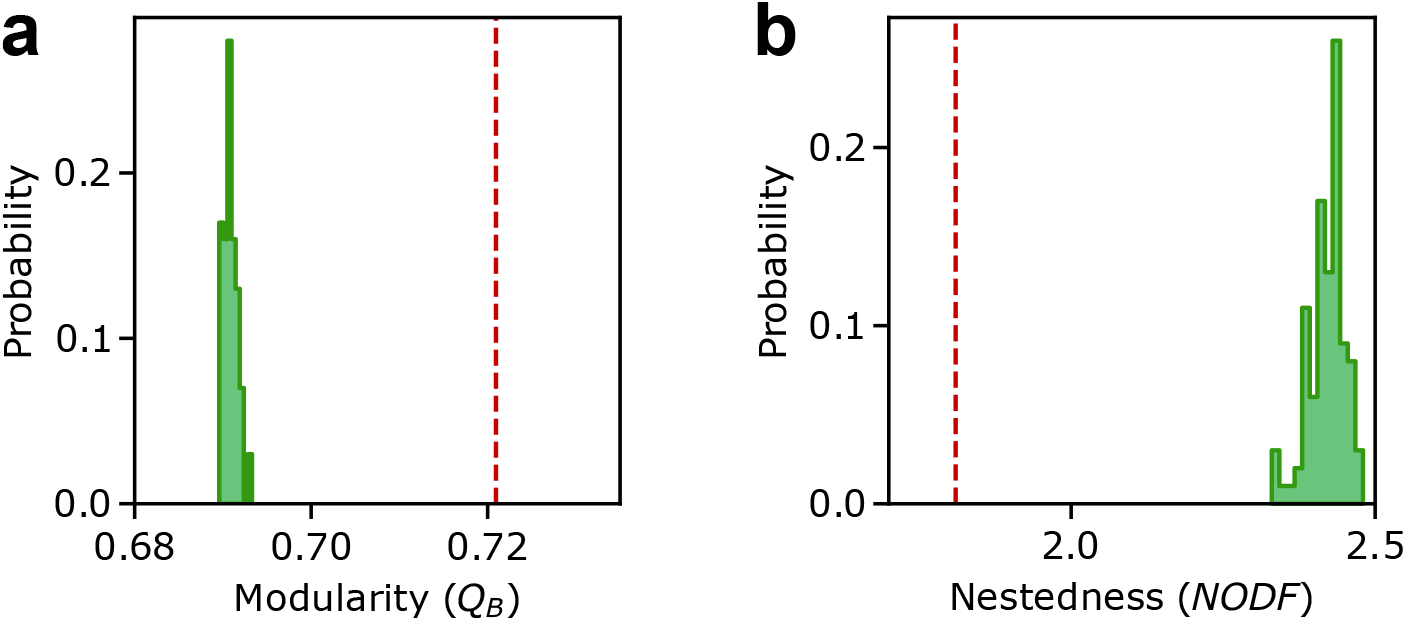
Nestedness and modularity in the photobiont-mycobiont network. Here we show (a) modularity denoted as *Q*_*B*_ and (b) nestedness calculated as the full network NODF (see Methods). The empirical values are shown as vertical red dashed lines, while bootstrap null model distributions are shown as a histogram of probability density obtained from 100 independent randomizations of the network.

### Differences between topological versus taxonomic signatures of guilds

Guild organization has been traditionally linked to photobiont identity (or “photobiont-mediated guild”), and more precisely, at the genus level^27,31,47^. Following this approach, Figure 4a shows the PMAN where species have been labelled according to the genus of their closest photobiont (using a label propagation algorithm, see Methods). We can see the labelled network comprises several contiguous clusters containing related species, i.e., belonging to the same genus. Here, the network appears to display a higher density of connections within each of the guilds than with outside nodes, which suggests that guilds display a common structural signature based on modularity.

**Figure 4.**
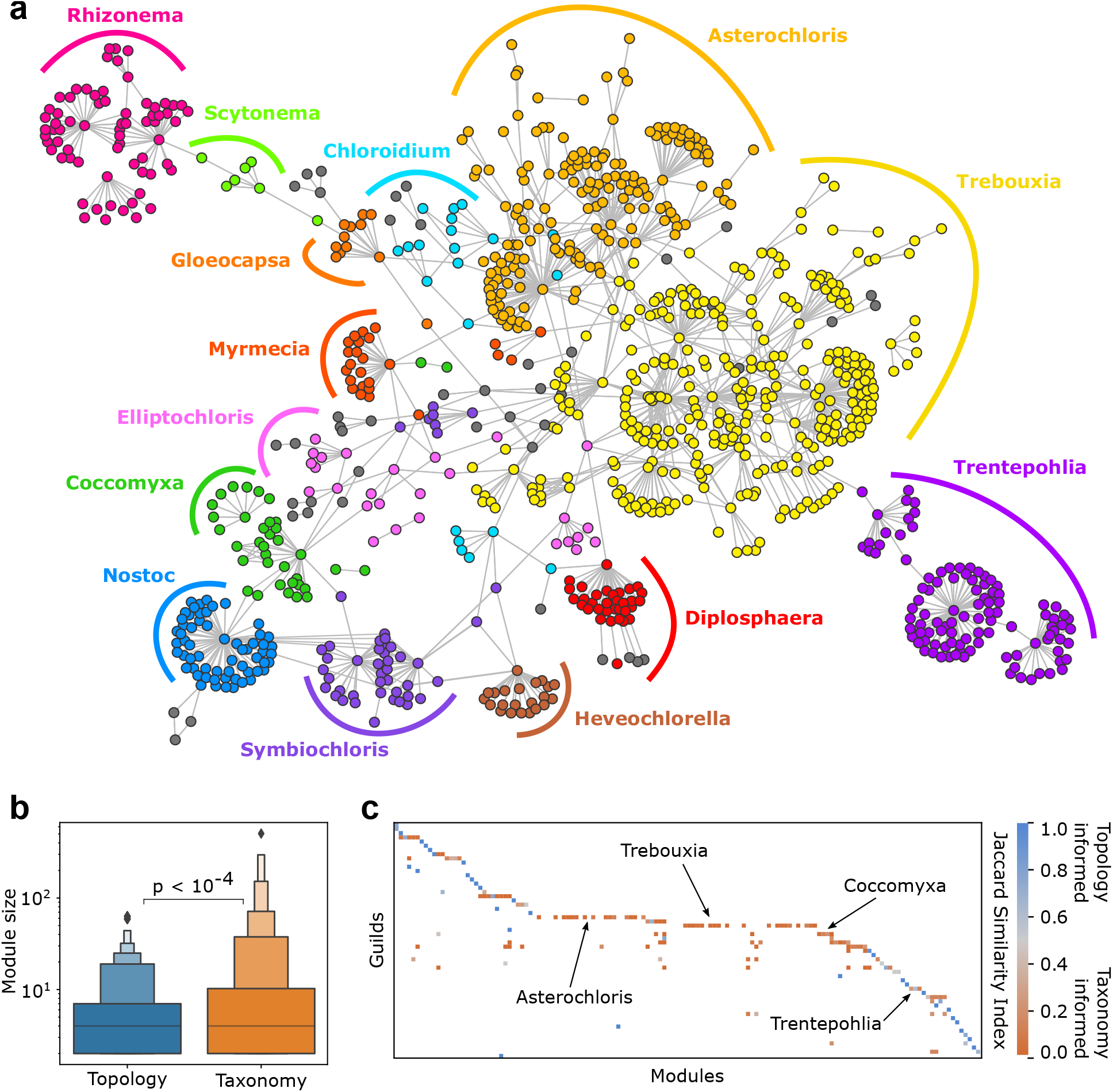
Comparing topological and taxonomic signatures of photobiont-mediated guilds. (a) Guilds predicted by photobiont taxonomic information in the largest subweb of the photobiont-mycobiont association network. Large guilds are represented by color, while minor guilds are represented by grey (clusters with fewer than five nodes). Network layout was automatically generated with the FMMM algorithm. (b) Module size distributions for two methods of guild allocation: a topological definition based on network modularity (blue) and a taxonomic definition of guild membership based on photobiont genera (orange). Taxonomic guilds are significantly larger than the topological modules (p-value obtained from t-test statistic). (c) Topological modules are incorporated into guilds. For each module-guild pair, we show the Jaccard’s similarity index, which is the intersection over union of the two sets of species. Some guilds (in blue) correspond to topologically defined modules, while others include several modules (in orange, see text).

We compare topological modules to well-known guilds to study the link between taxonomically-specified guilds and modularity (see Methods). In the PMAN there are 56 taxonomy-defined guilds and 140 topological modules. We find that topological modules are statistically smaller than the taxonomy-defined guilds, with an average size of 7.61 species per module versus 26.11 species per guild (*p <* 10^−4^, see Figure 4b). Smaller guilds have a strong matching with topology-predicted modules (as measured by Jaccard’s similarity^49^ between sets of species belonging to each community in Figure 4c), whereas larger guilds include several smaller modules embedded within them (see the highlighted cases of *Asterochloris* and *Trebouxia*). There are 21 guilds with average Jaccard similarity less than 0.5, indicating a significant discrepancy between topology and taxonomy, while the remaining 35 have very similar communities.

Comparison of taxonomic and topological information implies the existence of other structural patterns beyond the mesoscale defined by guilds. Empirical support to this hypothesis is found in the internal structural diversity of two major guilds: *Coccomyxa* and *Trebouxia*. These guilds are well established in the literature^27,30^ and provide a reliable microcosm whose individual components been extensively characterized, making them an ideal test-bed for our approach. Figure 5a shows a schematic representation of guilds proposed by Rikkinen^27^, while Figure 5b-c, on the other hand, shows the sub-networks of the same guilds with the most current information. These representations clearly show distinct characteristics, but the network approach allows us to measure these differences more precisely.

**Figure 5.**
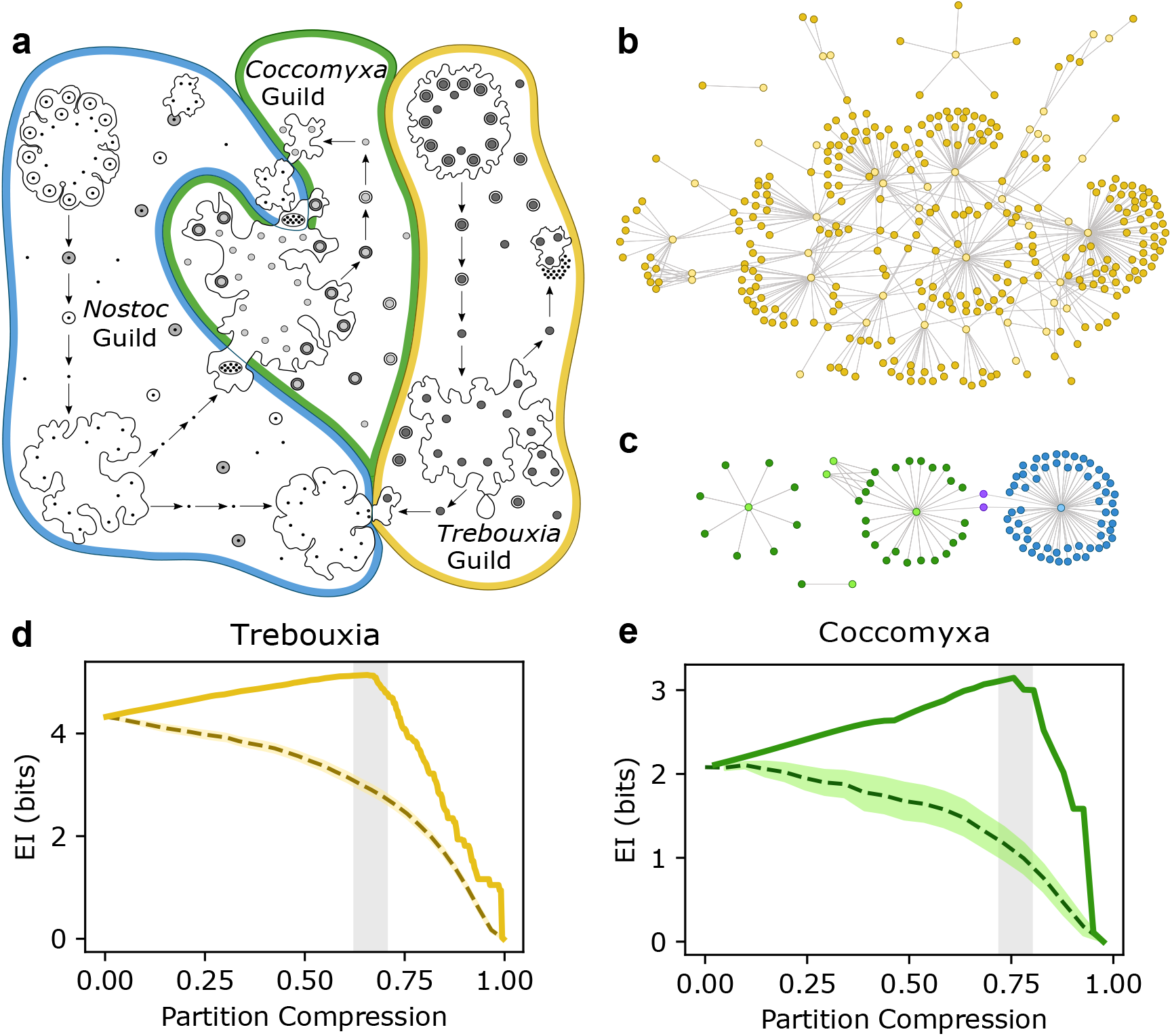
Effective information of photobiont-mediated guild networks. (a) Schematic representation of the guild relationships and composition as proposed by Rikkinen (adapted from^27^). An updated view of the *Trebouxia* guild (b), as well as the *Coccomyxa* guild (c). Guild internal structure can be quantified by its effective information (EI, in bits) at various degrees of compression (d-e, see methods). Here we show the effective information of random partitions for each guild (dashed line, 1000 independent random partitions of the guild graph at any given compression, shaded area in color encompasses one standard deviation of the sample) and the partition with the highest effective information for each compression value (solid line). (d) *Trebouxia* effective information suggests large amounts of internal structure that can be compressed into multiple (≈ 100) submodules within the guild. (e) *Coccomyxa* paints a similar picture, with its effective information across scales showing a single peak about 0.75 partition compression.

Figure 5d-e shows the effective information of the *Trebouxia* and *Coccomyxa* guilds using varying degrees of compression or aggregation into coarse-grained modules. Effective information^50^ evaluates if a coarse-grained representation of a network (i.e., nodes aggregated into modules) involves less uncertainty than a microscopic description of the same system (see Methods). For low levels of compression, there is limited aggregation of nodes into modules, and the *Trebouxia* guild shows the most intricate structure. Without exception, effective information increases with compression, meaning that aggregation into larger modules makes network representations less uncertain. This is followed by a peak effective information at a given scale (shown as a shaded grey region in Figure 5d-e). In both cases, the characteristic scale of maximal effective information at intermediate values suggests that these guilds are better represented as groups of distinct substructures rather than as monolithic communities. These results reinforce the predicted differences between taxonomy-defined guilds and topological modules.

### Guild interconnectivity matches major photobiont groups

At a bigger scale, it is important to assess whether there the interconnectivity among guilds is structured or not. Are there any non-trivial patterns beyond the level of guild organization? We build a guild interaction network (GIN) where nodes represent the major guilds (consisting of at least 5 species) and edges connect each pair of guilds that share at least one species (see Methods). Figure 6 depicts the adjacency matrix (a) and the circular layout (b) for the GIN. Using adjacency as a surrogate for distance in this network, we can estimate node similarity and cluster guilds based on topology using the fastcluster algorithm^51^. The photobiont phylogeny can be partially recovered in the GIN: several large green algae guilds cluster together (from *Heveochlorella* to *Asterochloris*) followed by smaller modules containing some green algae, *Xanthophytes* and cyanobacteria located in its own domain. In Figure 6b, the network representation allows us to show additional information: edge width is proportional to the number of shared species amongst guilds, while node size is proportional to the number of species in each guild. Guilds are colored according to photobiont type, green for green algae, blue for cyanobacteria and yellow for yellow-green algae. In agreement with our prior clustering analysis, every cyanobacterial guild groups together except for *Nostoc*. This implies that the large-scale organization of guilds is not random, but rather the result of biological traits or evolutionary constraints that bring physiologically similar species closer together in the network.

**Figure 6.**
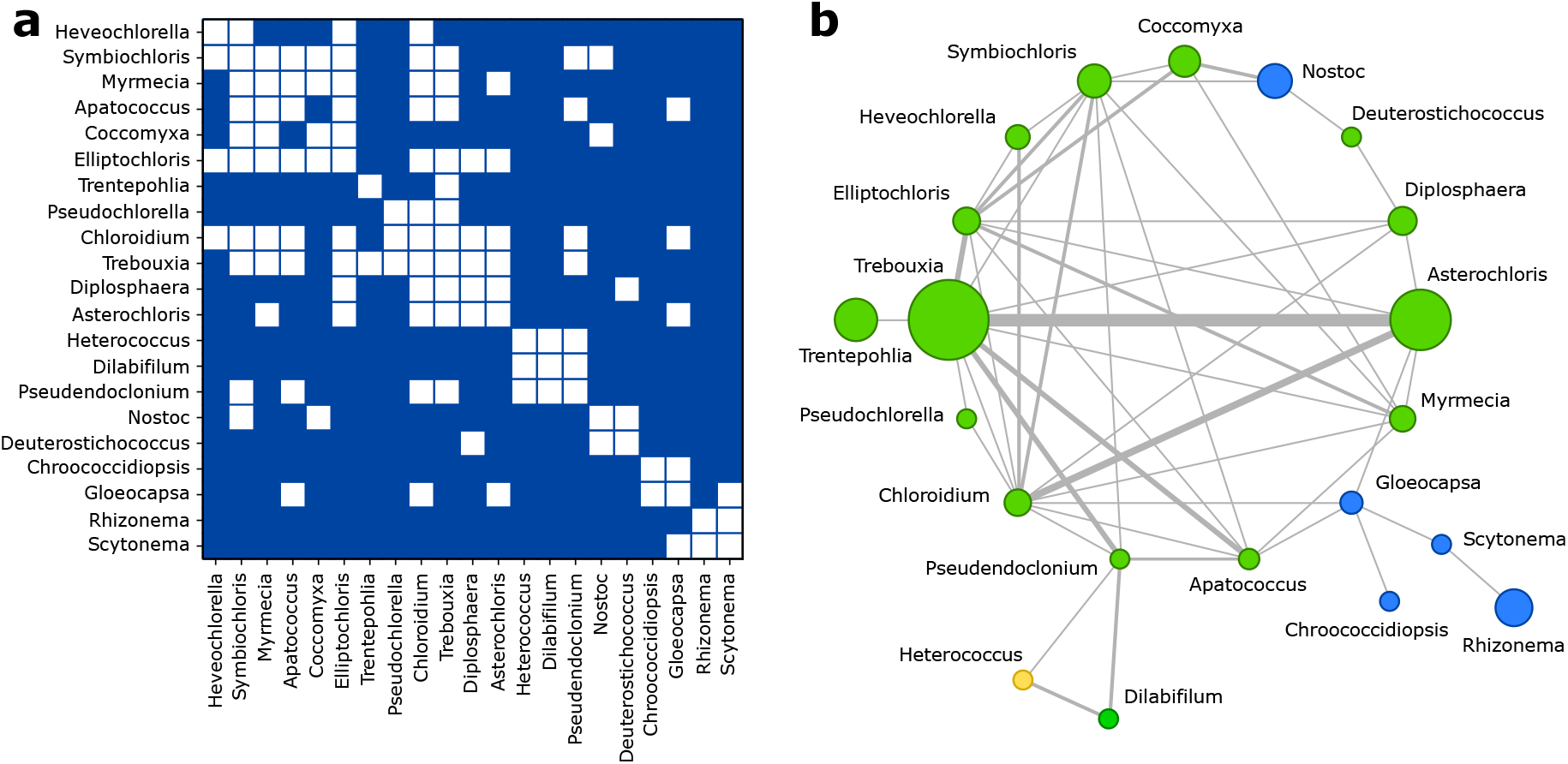
Global connectivity between photobiont genera. Using the photobiont genus as the foundation for lichen guilds, we reconstruct the guild network’s adjacency matrix (a), where white squares represent at least one mycobiont shared by photobiont genera. Photobionts are sorted using their pairwise distances and fastcluster algorithm (see Methods). In (b) we show the corresponding network representation, where link widths are proportional to the amount of shared mycobionts between photobiont genera, node sizes are proportional to the number of species belonging to that guild and and are colored according to their group: green for green algae (*Chlorophyta* and *Charophyta*), blue for cyanobacteria and yellow for yellow-green algae (*Xanthophyta*).

### Guild and species contribution to network robustness

We investigate the PMAN’s ability to withstand various types of perturbation while also providing some topologically-based insights to preservation efforts through the identification of keystone species and guilds. The degree of network fragmentation caused by species extinction is quantified using global efficiency, which reflects how costly it is to convey information across nodes. Global efficiency falls as the distance to a given node increases, eventually approaching zero in a fully disconnected system (see Methods).

Figure 7a shows the effects of different strategies of species removal on the global efficiency of the PMAN: random removal of species (blue), removal based on degree (green) and removal based on centrality (yellow). The PMAN is particularly resistant to random component failure; even after removing 120 nodes at random (about 10% of the network), the overall efficiency of the system is barely affected. This is consistent with many observations in heterogeneous biological networks^52^, which contain a backbone of nodes driving global connectivity, but most species have few connections and their extinction has a much more limited influence on the system’s structure. In contrast, the network is particularly vulnerable to targeted node removal, whether based on degree or node centrality. While topological network properties are unlikely to inform ecological damage, this sort of study might help guide conservation strategies that seek to prevent fragmentation and the buildup of ecological damage.

**Figure 7.**
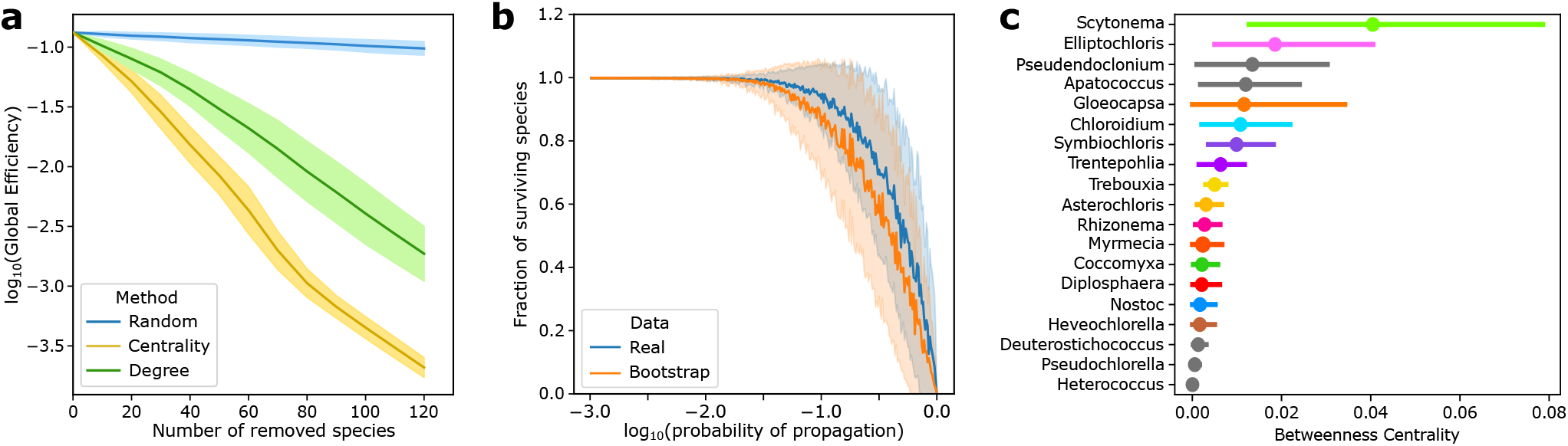
Guild and species contribution to network robustness. (a) Decay of global efficiency (and robustness) with increasing number of extinct species for different removal strategies. Centrality-based (yellow) removal of species affects the global path lengths and connectivity more than degree-based (green) or random removal of species (blue). (b) Surviving fraction of species in a cascading extinction event in the real (blue) and the bootstrapped (orange) data set. All simulations include 100 independent replicates in (a) and (b), with the solid lines representing the mean value of the distribution and the shaded area around it encompasses one standard deviation of the sample. In (c) we show the average betweenness centrality for the species in each guild, bars stand for the standard error in the distribution.

**Figure 8.**
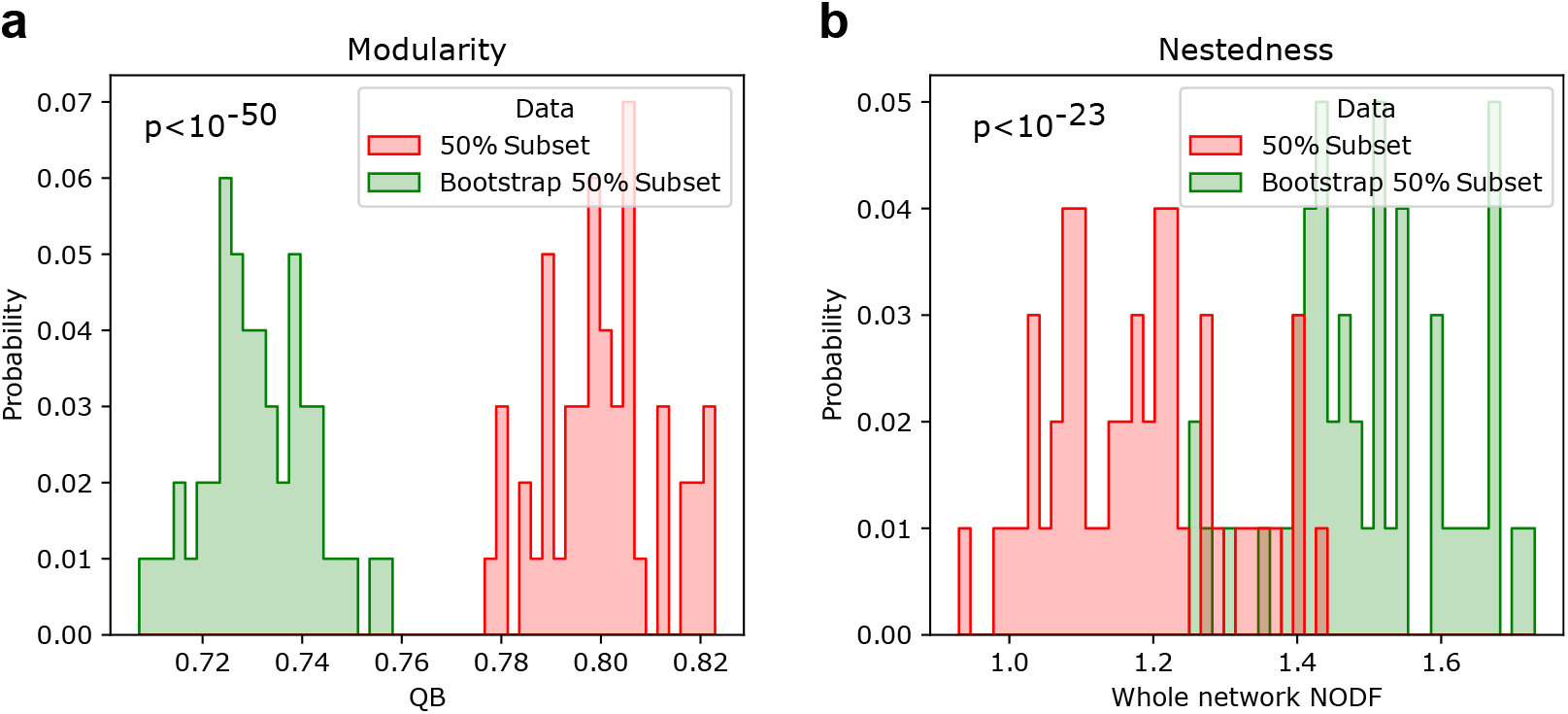
Validation of patterns in subsampled networks. Modularity (a) and Nestedness (b) patterns for subsampled networks in green (containing randomly half of the reported interactions) as well as their edge randomization counterparts (red). Distributions shown contain 50 independent data points of the subsampling and edge randomization each. The patterns reported for the full network of more modular than expected and less nested than expected are maintained, thus suggesting that these are not caused by sampling biases.

Figure 7b shows a simulated cascading extinction event, in which random species disappear and with a given probability this effect is propagated to neighboring species. When the chance of propagation increases, the amount of affected species increases, reaching the totality of the system when the probability is 1. This type of analysis is carried as a comparison between the the real data set (blue) versus an ensemble of bootstrapped networks (orange) acting as negative controls. On average, the real data set is more robust to cascading extinctions for a wide range of values. The modular pattern present in the PMAN (which is removed in a bootstrap) acts as a firewall, limiting the reach of harmful effects to local domains.

Some guilds and species have a greater influence than others in providing global resilience due to their placement within the network as well as their local characteristics (Figure 7c). Of particular interest are the guilds *Scytonema* and *Gloeocapsa*, which act as bridges to the major cyanobacterial guild *Rhizonema*. In addition, species within the major guilds *Elliptochloris, Symbiochloris* and *Chloroidium* have relatively high centralities. Remarkably, these are fragmented guilds (they do not form continuous clusters in Figure 4a) and their constituent species often lie at the intersection between other major guilds in the network.

## Discussion

At the core of the lichen symbiosis is the association between two unequal partners, a fungi and a phototroph. The flexibility of this association is key to understand the resilience of lichen species to polluted and antropogenic ecosystems^10^ and their capacity to adapt to new environments^6^. Understanding and documenting single partnerships has been a major focus of lichen research. However, mycobiont-photobiont partnerships do not happen in isolation, they are part of a larger web of interactions that shapes their chances of formation and success. While much effort has been invested in characterizing individual associations, the global network of interactions that maintain the lichen symbiosis remains unknown. Using the most comprehensive data set of mycobiont-photobiont partnerships to date, we addressed the multi-scale nature of lichen symbiosis from a network perspective.

Ecological networks have emerged as a powerful tool to formalize biotic interactions, including mutualistic, antagonistic and more complex relations^53^. This characteristic makes them particularly useful to study the so called photobiont-mediated guilds^27^: communities of lichens that share a common set of photobionts and facilitate each other’s propagation, but also compete for space and resources. Lichens depend on a set of coupled ecological interactions involving species dispersal, facilitation and competition, which are in turn shaped by guild structure^30^. Mycobionts within a guild are able to promote the establishment of one another into new habitats by extending the ecological range of their partners^31^. For example, spore-producing lichens that require a compatible photobiont upon reproduction are indirectly facilitated by the asexual lichens within their guild already established in that environment^28^.

Previously, guilds have been defined by photobiont taxonomic identity, but this characterisation has been constrained by the scope of the data sets, which have often included a limited amount of species or geographical locations^16,21,30–34^. An aggregated network perspective allows us to tackle the structural signature of lichen guilds and provides a solid foundation to address how the general patterns of association in the lichen symbiont network affect its robustness.

As communities of species, guilds occupy a mesoscale in the photobiont-mycobiont association network. By sharing a common set of photobionts, species within a guild should interact more frequently with other species in it than with those outside. In network theory terms, this relates to a common structural pattern of clusters of strongly interacting species (i.e. high modularity). Theoretical and empirical studies predict that in general many ecological communities are composed of several loosely interrelated compartments, each containing multiple species^54^. Recent studies involving small sets of lichen species and their interactions have revealed diverging structural patterns: from distinct clusters of strongly interacting species in *Peltigera* lichens^36^ to an embedded specialist-generalist (nested) structure in *Nephroma*^31^. Different explanatory causes have been proposed for the observed nested and modular patterns, including evolutionary constraints^36^ as well as the variations in the nature of the underlying ecological interactions^31^ (i.e. mutualistic vs. non-mutualistic relations). Analysing the full set of interactions compiled from past research is essential in determining whether the reported structure of the PMAN are validated or the result of limited data.

Our analysis shows that the PMAN displays a statistically significant modular and non-nested organisation when compared to the null model. This is consistent with previous observations of *Peltigera* lichens^36^. Furthermore, we compare the processes of guild allocation based on photobiont taxa and topologically defined guilds, testing pre-established assumptions regarding guild structure and composition. Beyond the proposed guild mesoscale, we can identify other relevant scales in lichen symbiosis. Small taxonomy-defined guilds usually match with topological modules, and are typically composed by a single photobiont linked to a few mycobionts. Instead, larger guilds like *Asterochloris* and *Trebouxia* are rather complex communities with many interacting sub-modules. These guilds are characterized by a peak effective information at intermediate compression values, highlighting a good match between the effective information and modularity approaches to community detection. This newly reported scale in the PMAN suggests that additional underlying ecological or generative constraints (such as habitat range or trait-dependent associations) may play an important role in shaping guild structure.

According to the GIN analysis, trait-dependent associations may also inform large-scale interactions between photobiont genera. We found that guilds are connected in a non-trivial manner consistent with the evolutionary history of photobionts, separating algae and cyanobacteria into two identifiable clusters. A possible explanation to understand these findings might be the unique way by which some cyanobacteria are lichenized. Lichens harboring cyanobacterial and algal partners segregate them in the same organism using specific organs called cephalodia^55^. The use of specific and non-universal morphological structures for cyanobacterial symbionts points at a distinct proximal evolutionary origin for these innovations and highlights the importance of morphological novelty as a driver of the patterns observed in the GIN. Putting these two structural patterns together, the PMAN recapitulates evolutionary history at the largest scales, but is inconsistent with taxon-based guild definitions at the smallest scales.

The network approach has the potential to give insights into the underlying symbiotic associations. The nature of the lichen symbiosis, whether mutualistic, antagonistic, or somewhere in between, is a subject of debate within the community^56–59^. Some authors regard lichen symbiosis as mutually beneficial among partners^34,59^, while others propose a more complex relationship analogous to photobiont domestication by mycobionts^58,60^. Under the lens of facultative mutualism, symbiosis can be malleable, defined by additional factors like species density^61^ or the quality of the external environment^57^. In harsh conditions, photobionts in association with fungi are better protected from external challenges like predation, UV radiation and extreme temperatures^62^. Yet in optimal environmental conditions there are opportunity costs to living in association; the photobiont might be better off free-living instead of transferring metabolites to its host in exchange for a protection that is not required.

Theoretical studies have used different arguments to explain the presence (or absence) of modularity and nestedness in ecological networks, including ecological stability, spatial constraints, or the strength of coevolution^63–65^. Evolutionary network models have shown that the topological features of an ecological network determine its resilience, stabilizing structural patterns based on the types of interactions it shows. For instance, nestedness is often the key pattern found in mutualistic networks^38,63^, while modular structures are commonly associated with antagonistic networks^43,63,64^. Modularity and nestedness are negatively correlated but they can also co-occur in sparsely connected random networks^44^. This observation highlights the remarkable structure of the PMAN, which even at low connectance values displays modular, non-nested structures. An important aspect is that nestedness can also be produced for-free by the evolutionary generative rules^66^, and it is still unclear if nestedness correlates with stability according to more recent metrics^67^. Nestedness and modularity are not exclusive to particular interaction types^68^, and they can also coexist, occupying different scales in the ecological network (e.g. modules that are themselves nested^39,69,70^) or different dimensions of hypergraph ecologies^71^. Our results suggest the presence of non-mutualistic interactions among lichen symbionts, whether antagonism or facultative mutualism^43,63,64^.

Network structure can have a significant impact on the resilience and persistence of species and ecosystems^52,72^. How perturbations propagate in ecological networks has been the subject of numerous studies, bridging the gap between topology and population dynamics. Fragmentation of the ecological network can have a detrimental influence on effective diversity and disrupt the flows of matter, energy, and ecosystem services^73,74^. In turn, diversity has a buffering effect in accumulation of damage and the effects of perturbations in ecological systems^75^. However, not all species have the same contribution to network resilience. Removal of central or highly connected species can pose larger threats to network integrity (see Figure 7c). Modular structure also impacts the spread of cascading extinctions, confining perturbations to small groups of species^54,64^. The symbiotic network presented here benefits from both of these patterns: hub species keep the network connected even when individual species are randomly removed, and the highly modular structure increases resilience by functioning as a firewall compartmentalizing damage.

In conclusion, we found evidence of the topological signature of guilds in the dataset of aggregated photobiont-mycobiont associations. The presence of a distinctive scale in the effective information reveals a gap between taxonomically defined guilds and topological modules. This stands in stark contrast to the largest patterns of interaction in the network, which closely follow major evolutionary groups. Our analysis shows that some guilds have a greater influence than others in terms of network robustness. Future work will have to address the biological reasons explaining the origin of the reported structural patterns in symbiont networks. Incorporation of topological patterns in theoretical models will improve our knowledge about network robustness, which is essential to anticipate biodiversity losses and extinction cascades.

## 1 Materials and methods

### 1.1 Data set

To build the bipartite network of photobiont-mycobiont associations, we use the data from a recent meta-study^37^. This work compiles over 200 publications in the field of lichen symbiosis, describing natural interactions between symbionts. The data was supplemented with additional well-interactions (Lobaria’s cyanobiont partners) that were missing in the original data set. The meta-study by Sanders and Matsumoto focuses on publications after 1988 Tschermak-Woess’ influential review and especially those publications providing molecular evidence of species identity. This dataset is, however, biased towards geographical locations in the western world (namely, Europe and North America). Most symbiont pairs are genetically validated through sequencing efforts and correspond to a recent wave of publication in the field, but the earliest publications in the data set (e.g. “A monography on algal culture” by Chodat, 1913) do not provide the same degree of validation. Thus, the data set contains some degenerate information in which, symbionts are identified only up to the genus level instead of the species level (≈ 30% of the data). Similar patterns as those reported here were found in the analyses pertaining the non-degenerate network, as well as those limited to the largest subweb. From the listing of naturally observed associations, we can define a photobiont-mycobiont association network (PMAN) *G* = (*P, M, E*) in terms of the two disjoint sets of photobiont *P* and mycobiont *M* species and another set of edges (*i, j*) ∈ *E* capturing the symbiotic association between partnering species *i* ∈ *P* and *j* ∈ *M*. The graph *G* can be also written as a biadjacency matrix *B*_*i j*_ with dimensions corresponding to the size of each set of species. Any given entry in the matrix equals zero if there is no interaction between species *i* and *j* and equals 1 otherwise. From the PMAN, we can further define a guild interaction network (GIN), Γ = (*V, I*) composed of a set of guilds *u, v* ∈ *V* and their interactions *I*. Links (*u, v*) ∈ *I* in this network indicate whether a pair of photobionts belonging to different guilds share one or more mycobionts. We can compute the (weighted) adjacency matrix *A* = [*A*_*uv*_] of the GIN as follows:

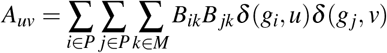

where the membership function *g*_*i*_ indicates what guild the photobionts species *i* belongs to, and *δ* (*a, b*) = 1 when *a* = *b* or 0, otherwise. The *V* × *V* square matrix *A* represents an unipartite network whose entries measure how many mycobionts are shared between pairs of photobiont genera.

#### 1.1.1 Taxonomical definition of guilds using label propagation

In order to establish taxonomical guilds we use a variation on the classical label propagation algorithm by Raghavan, Albert and Kumara^76^. In our bipartite label propagation algorithm, photobiont nodes are labeled with their genus, while mycobionts are left unlabeled. Mycobiont nodes then acquire their surrounding photobiont tags sequentially but in a random sequence, and each community grows isotropically. In the case that a mycobiont has neighbors in different communities, a dual community allocation is proposed for that given node. At the end of this iterative process, nodes tagged with the same labels are grouped together as guilds. This is a simple and effective method for discovering any underlying structure in the PMAN network that is led by taxonomic information.

### 1.2 Network metrics

#### 1.2.1 Modularity

A number of metrics have been used to assess the modular structure of ecological systems, but a popular method examines the degree of modularity at the topological level^77^. Given a partition of the set of nodes given by the function *c*(*i*) indicating the module label to which the node *i* belongs, modularity *Q* calculates the difference between the proportion of links inside each module and the predicted fraction when connections are randomly rewired:

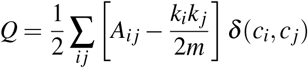

where *A* = [*A*_*i j*_] is the adjacency matrix of the (unipartite) network, 2*m* = ∑_*i j*_ *A*_*i j*_ is twice the number of edges, *k*_*i*_ = ∑ _*j*_ *A*_*i j*_ is the degree of node *i*, and *δ* (*a, b*) = 1 when *a* = *b* or 0, otherwise. The previous definition must be modified for a bipartite network to add the requirement that no connections exist between nodes of the same type^78^. In this case, the bipartite modularity (*Q*_*B*_) is defined as:

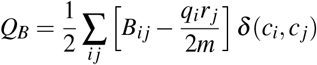

where *B* = [*B*_*i j*_] is the bipartite adjacency matrix, and *q*_*i*_ = ∑ _*j*_ *B*_*i j*_ and *r* _*j*_ = ∑_*i*_ *B*_*i j*_ is the degree of photobionts and mycobionts species, respectively. Here, we measured the bipartite modularity of the PMAN using a custom implementation of the biLOUVAIN algorithm^79^ in Python 3.9, which heuristically finds a modularity maximizing arrangement of photobionts and mycobionts into a non-predetermined number of modules. Initially, each node is allocated to its own community and then, iteratively, the algorithm finds and applies a community allocation change (swapping the module of a single node) that yields an increase in the overall modularity. This process stops when the algorithm cannot find a community swap that further increases the global modularity beyond a predetermined threshold *ρ* (in our case *ρ* = 10^−6^). The bipartite modularity *Q*_*B*_ assesses how often a particular annotation of nodes into modules corresponds to interactions that are mostly inside each module (*Q*_*B*_ = 1) versus mostly outside of each module (*Q*_*B*_ = −1).

#### 1.2.2 Nestedness

Nestedness is a global measure of the propensity of low-degree species to interact with a subset of highly interconnected species. This pattern is clearly identified by the network architecture; nestedness is a systematic arrangement of non-zero entries in the adjacency matrix. We compute the nestedness using the overlap and declining fill (NODF^80^) for the bipartite graph *G* = (*P, M, E*) with biadjacency matrix *B* = [*B*_*i, j*_]:

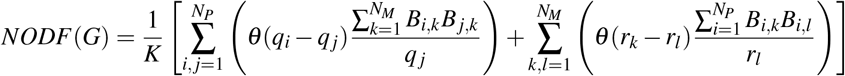

where *K* = [*N*_*P*_(*N*_*P*_ − 1) + *N*_*M*_(*N*_*M*_ − 1)]*/*200 is a normalisation constant, *N*_*P*_ = |*P*|, *N*_*M*_ = |*M*|, *θ* is the Heaviside function with *θ* (0) = 0, and *q*_*i*_ and *r* _*j*_ are the degrees of nodes *i* ∈ *P* and *j* ∈ *M*, respectively. A high NODF value implies that certain species’ interactions are a subset of other generalist species’ interactions (and so the network demonstrates nesting), whereas a low value indicates clustering (which is consistent with high modularity). After determining the empirical nestedness values, they are compared against null model predictions to assess their statistical significance. Nestedness is sensitive to network size *N* = |*P*| + |*M*| as well as network connectivity, according to theoretical and empirical investigations. We must compare empirical data with a bootstrap technique that keeps the size and fill of the PMAN’s adjacency matrix since we do not know the underlying null distribution of the test statistics.

#### 1.2.3 Global efficiency

Latora and Marchiori^81^ introduced a network metric of global integration that is inversely proportional to the average path length. The mean global efficiency (*E*(*G*)) of a network *G* can be defined as follows:

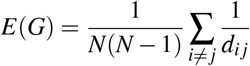

where *d*_*i j*_ is the length of the shortest path connecting any pair of nodes *i* and *j*, and *N* is the network size. Distances between pairs of nodes in disjointed subweb are infinite, as there is no path that can reach from the original node to the destination. An advantage of using the inverse of path length (instead of the alternative average distance) is that it allows computation of a finite efficiency value for graphs with fragmented networks. Efficiency calculations were carried out using the NetworkX version 2.8 of the algorithm for Python^82^.

#### 1.2.4 Betweenness centrality

Centrality measures propose a method to evaluate node or link importance given a certain network topology. Betweenness centrality (BC) of a given node is the fraction of shortest paths that pass through that node, considered over all other possible pairs of nodes within the network. A low betweenness centrality (*BC* = 0) means that the node is not in the path to reach other nodes, and thus has no effect in the flow of information within the network. High Betweenness centrality (*BC* = 1) means that the node is always involved in the traffic of information in the network and its removal could have major impacts in the network’s information flow, whether through rerouting of shortest paths (making them longer and thus a less well connected network) or simply fragmentation impeding further communication between sets of disjointed nodes. Here, a bipartite or unipartite implementation is essentially the same, but a consideration must be given to the sampling mechanism of node pairs. The process can be substantially sped up at the cost of reliability by reducing the fraction of pairs considered in the analysis. All calculations shown in this publication correspond to the full set of nodes, computed following the standard algorithm by Borgatti and Halgin^83^, as implemented in NetworkX version 2.8^82^.

#### 1.2.5 Effective information

Effective information (EI) is a recent metric proposed by Klein and Hoel^50^ that quantifies the information contained in the topology of a network and can be used to signal the presence of higher informative scales. EI can be expressed as the trade-off between the determinism and degeneracy associated with a given topological configuration:

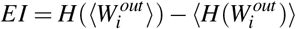

where the first term corresponds to the degree of determinism or certainty associated to a random walker traversing the network in every node, and the second term captures the system’s degeneracy or entropy in weight distributions. Here, *H*(*X*) = −∑_*n*_ *P*(*x*_*i*_) log_2_ *P*(*x*_*i*_) represents the Shannon’s entropy of a given distribution *X*, and the distribution 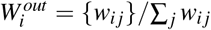 is the normalized set of weights from node *i* to every other node *j*. These weights can be understood as the distribution of probabilities a random walker might take when visiting node *i*.

Klein and Hoel expanded the above formalism to an algorithmic procedure that detects higher informative scales in network structures. Formally, these scales are identified with coarse-grained representations of the system in which certain nodes have been aggregated together, causing the system’s effective information to increase. During the aggregation process, the network is mapped into a Markovian chain, with outgoing connections from any macro node corresponding to conditional probabilities of random walkers in the original, unaggregated network. Finding informative scales entails iteratively proposing new macro nodes, which are assemblages of existing micro nodes or a mixture of micro and macro nodes. The algorithm greedily aggregates nodes in order that maximize the effective information of the resultant network at each step, a process known as causal emergence. When no aggregations to improve effective information beyond a given threshold are detected, the procedure ends (threshold used *ρ* = 10^−4^). We extended the above approach by determining the minimal descent from the maximal effective information configuration. Beginning with this peak, we try random aggregation steps while retaining any partitions that involve a minimal loss in effective information. Compression associated with a given partition *P*_*t*_ (e.g. in Figure 5) is *C*(*P*_*t*_) = 1 − *N*_*t*_*/N*_0_, where *N*_0_, *N*_*t*_ are the initial number of nodes and nodes in partition after aggregation *P*_*t*_, respectively. The base code library used is einet 1.0.0 for Python^84^.

### 1.3 Robustness to species loss

We tested the effects of species removal on the global efficiency using three different strategies for choosing nodes: (1) Random, (2) Degree-based, and (3) Centrality-based. For the two latter strategies node choice was carried out using a propensity vector incorporating the normalized degree sequence and the normalized betweenness centrality (calculated as described in the prior section).

We also analyzed the effects of species extinction propagation in the symbiont network through a simulation of extinction cascades^85,86^. In these simulations, a node is randomly chosen from the network (both in the empirical data set and the bootstrapped data set) and an “infection” process is carried out. While a node is infected (or becoming extinct) it can affect neighboring nodes with a given probability (a parameter fixed for each simulation). If new nodes become extinct then the process continues until no new extinction events are produced. The reported results are the fraction of surviving species at the end of the extinction cascade. These simulations are carried out in 100 independent replicates (meaning a replicate for 100 different bootstraps of the original data set for the case of the negative control).

### 1.4 Statistical validation

#### 1.4.1 Statistical significance of modularity and nestedness under sampling biases

Following standard practices in studies of symbiotic networks^48^, we further validated the patterns reported here by performing similar analyses on randomized subsets including only half the documented interactions. We compared a set of 50 subsampled networks with the same amount of bootstrap edge randomization of these subsampled networks. The subsampled networks containing 50% of the total interactions were more modular and less nested that their bootstrap counterparts, reinforcing the perspective that the patterns reported here are meaningful and unlikely to be caused by limited sampling of the underlying interactions. Significance of differences in these metrics was quantified with a *scipy* t-test statistic, yielding the significance p-values of *p <* 10^−5^ for both modularity and nestedness.

#### 1.4.2 Null models of bipartite networks

Validating the statistical significance of structural patterns is a common approach in network studies (i.e., two classic examples are motif analysis^87^ and community detection^88^). The goal here is to determine which empirical patterns depart from a baseline null model in order to give evidence that these patterns are a feature of the system under study rather than a result of other qualities. In this paper, we employ a bootstrap randomization of the bipartite network which fully preserves the original degree distribution^89^. Interactions are allocated using a propensity vector for node identity that takes into account the original node degree in each compartment^42^. Using this strategy degree heterogeneity is preserved (as well as any network properties stemming from it), but node-node correlations are removed from the system.

## Acknowledgements

We thank Julia Adams, Blai Vidiella and Josep Sardanyés for helpful discussions on the manuscript. SV is supported by Grant PID2020-117822GB-I00 funded by MCIN/AEI/10.13039/501100011033. SDN is supported by the Beatriu de Pinós postdoctoral programme, from the Office of the General Secretary of Research and Universities and the Ministry of Research and Univertisites (2019 BP 00296) and the support of the Marie Sklodowska-Curie COFUND (BP3 contract no. 801370) of the H2020 programme.

## Author contributions statement

S.D-N. conceived the research, S.D-N. and S.V conducted the experiments, S.V. and S.D-N. analysed the results. All authors wrote and reviewed the manuscript.

